# Hidden Complexity of Pediatric Platelet Disorders: Functional Diversity and Unexpected Hypercoagulable Phenotypes

**DOI:** 10.64898/2026.05.27.728206

**Authors:** Taisia O. Shepeliuk, Evgeniya Melnikova, Praharsha Konde, Ekhson Holmuhamedov, Fazly Ataullakhanov, Michele P. Lambert, Ekaterina L. Grishchuk

## Abstract

Pediatric platelet disorders are commonly classified according to specific structural or functional abnormalities, yet it remains unclear how well these diagnoses capture overall hemostatic phenotype. Here, we combined quantitative single-cell platelet measurements with spatially resolved plasma clotting analysis to characterize pediatric patients with dense granule deficiency, platelet function defects, immune thrombocytopenia, and other inherited platelet disorders. Quantitative fluorescence microscopy revealed reduced dense granule abundance not only in dense granule deficiency but also in several patients from other diagnostic groups. Measurements of platelet adhesion, spreading, and calcium signaling identified substantial functional diversity, with individual patients exhibiting distinct combinations of abnormalities that were not predicted by diagnostic category. Unexpectedly, plasma clotting analysis frequently revealed hypercoagulable behavior, including accelerated fibrin clot growth and spontaneous fibrin formation, despite clinical diagnoses associated with platelet-related bleeding disorders. Hypercoagulable phenotypes occurred across multiple diagnostic groups and did not show a simple relationship with platelet functional abnormalities. Together, these findings reveal previously unrecognized complexity in pediatric platelet disorders and suggest that platelet and plasma pathways contribute independently to hemostatic variability. These findings argue that pediatric platelet disorders are best viewed as multidimensional functional phenotypes rather than isolated platelet defects and motivate broader integration of platelet and coagulation measurements in future studies.

## Introduction

When a blood vessel is injured, circulating blood cells and soluble plasma factors rapidly assemble into a clot that limits blood loss while maintaining vessel patency. This process requires platelets to adhere, activate, secrete bioactive mediators, and provide catalytic surfaces, while plasma coagulation reactions generate thrombin and fibrin that stabilize the developing clot and support its remodeling^1^. Although pediatric platelet-related bleeding disorders are uncommon, they encompass a diverse group of acquired and inherited conditions whose clinical manifestations often cannot be explained solely by platelet count or by a single laboratory abnormality. Many pediatric diagnoses are defined by limited functional assays or by specific structural defects, yet the extent to which multiple aspects of platelet behavior are altered within individual patients remains poorly understood^2,3^. As a result, biologically meaningful functional heterogeneity may remain obscured within clinically defined diagnostic categories.

Dense granule deficiency (DGD, δ-storage pool deficiency) illustrates these challenges particularly well. DGD is characterized by reduced dense granule (DG) number, content, or secretion and is among the most frequently reported inherited platelet disorders. Diagnostic evaluation often begins with platelet aggregation studies, yet standard aggregometry may fail to identify a substantial fraction of affected patients^4,5^. Electron microscopy (EM) therefore remains the principal method for direct assessment of platelet DGs^6^. However, EM-based diagnosis requires specialized expertise, examines relatively small numbers of platelets, and relies on laboratory-specific thresholds and analytical approaches. Consequently, reported prevalence, diagnostic criteria, and clinical associations vary substantially between studies. Importantly, DGD is typically evaluated through granule counts alone, whereas other potentially informative platelet phenotypes, including adhesion, spreading, activation signaling, and coagulation-related functions, have received considerably less attention^7,8^. Thus, DGD provides both a clinically relevant disease cohort and an opportunity to examine whether quantitative single-cell measurements can reveal functional features that are not captured by conventional classification approaches.

To address this challenge, we selected platelet phenotypes that capture distinct aspects of platelet biology - DG abundance, adhesion/spreading behavior, and calcium-dependent activation - and combined these measurements with spatially resolved plasma coagulation analysis. Our goal was to determine whether clinically defined pediatric platelet disorders are associated with characteristic functional signatures or whether substantial variability exists both within and across diagnostic categories. To assay platelet function, we applied a microfluidic imaging platform that enables quantitative analysis of individual platelets under controlled flow conditions (Figure 1A). Unlike conventional bulk assays, this approach provides access not only to population averages but also to cell-to-cell variability and distributional features within each sample. We quantified DG abundance using mepacrine fluorescence and simultaneously examined platelet adhesion, spreading, and intracellular calcium signaling, a dynamic indicator of platelet activation. Previous studies have demonstrated the utility of microfluidic approaches for investigating platelet behavior^9^, evaluating therapeutic interventions^10–13^, and studying hemostatic abnormalities^14–20^. However, comparable multidimensional analyses remain uncommon in pediatric patient cohorts.

**Figure 1.**
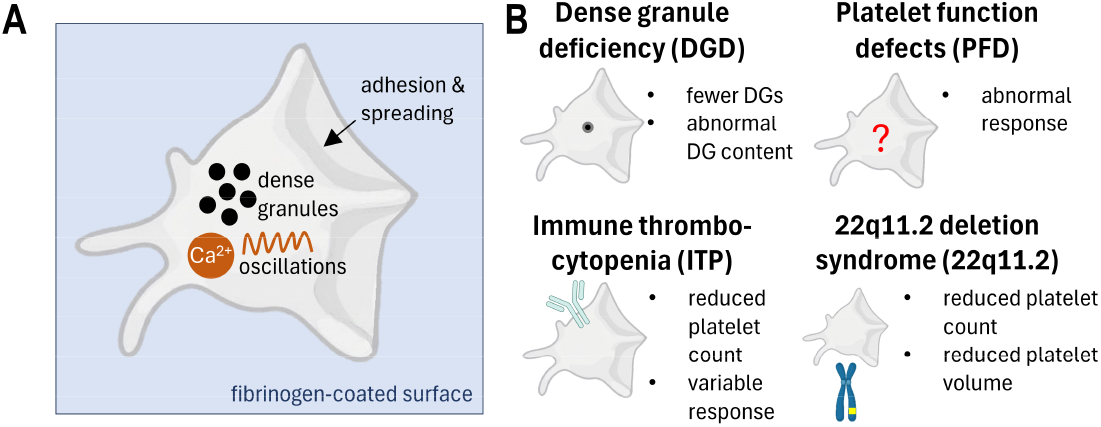
Overview of platelet functional assays and pediatric patient groups analyzed in this study. A. Schematic of single-platelet analysis on a fibrinogen-coated surface. Platelet adhesion and spreading, dense granule (DG) content, and intracellular Ca^2+^ oscillations were quantified as functional readouts. B. Pediatric patient groups included in the study. These groups represent distinct clinical contexts associated with altered platelet number, platelet volume, DG content, or platelet functional responses.

To determine whether observations made in DGD extend to other pediatric platelet disorders, we also analyzed several additional patient groups representing distinct biological mechanisms (Figure 1B). Platelet function defects (PFDs) comprise a heterogeneous collection of disorders defined operationally by abnormal platelet function testing rather than by a single molecular cause^21,22^. Depending on the underlying defect, abnormalities may involve signaling, secretion, aggregation, adhesion, spreading, or procoagulant activity. Immune thrombocytopenia (ITP) represents a fundamentally different condition in which autoimmune platelet destruction is accompanied by alterations in platelet activation state^23^. Despite similar platelet counts, patients with ITP often exhibit markedly different bleeding phenotypes^24,25^, suggesting substantial functional diversity among circulating platelets^26–29^. We additionally included patients with 22q11.2 deletion syndrome, in whom platelet abnormalities have been linked to haploinsufficiency of genes involved in megakaryocyte and platelet biology, including GPIBB^30^. Prior studies have reported thrombocytopenia, enlarged platelets, altered aggregation responses, and in some cases apparently normal platelet function, leaving the overall functional phenotype incompletely defined^31–34^. Finally, samples from patients with MYH9-related disorder and Gray platelet syndrome were included as biological reference cases with known platelet abnormalities affecting platelet structure and granule biology. Collectively, these groups provided an opportunity to examine whether different diagnostic categories correspond to distinct platelet phenotypes or whether functional variability extends across conventional disease boundaries.

Platelet-dependent pathways represent only one component of hemostasis. Plasma coagulation contributes independently to clot formation through the generation of thrombin and fibrin, yet platelet function and plasma coagulation are typically evaluated separately. Consequently, it remains unclear whether abnormalities in platelet behavior are accompanied by corresponding changes in coagulation potential or whether these systems vary independently within individual patients. To address this gap, we complemented platelet phenotyping with analysis of platelet-free plasma using the Thrombodynamics assay. Unlike conventional global coagulation tests, Thrombodynamics initiates clotting from a localized tissue factor source and quantifies spatial clot propagation, providing sensitive measurements of clot growth kinetics and spontaneous coagulation activity while requiring only small plasma volumes^35^.

Using this integrated approach, we examined DG abundance, adhesion, spreading, calcium signaling, and plasma coagulation in pediatric patients with DGD and other platelet-related disorders. We found that functional abnormalities frequently extended beyond the features traditionally associated with a given diagnosis and that individual patients often exhibited distinct combinations of platelet phenotypes. Unexpectedly, enhanced plasma coagulation was common across several diagnostic groups despite the presence of platelet functional abnormalities. Together, these findings reveal substantial functional diversity within pediatric platelet disorders and highlight the value of multidimensional phenotyping approaches that extend beyond conventional diagnostic classification.

## Materials and Methods

### Human ethics

Collection of pediatric human blood was approved by the Children Hospital of Philadelphia Institutional Review Board (№ 12-009829 and 08-006082). Collection of adult human blood was approved by the University of Pennsylvania Institutional Review Board (№ 855449). Written informed consent was obtained from adult donors and from parents or legal guardians of pediatric participants, with assent obtained when appropriate, in accordance with the Declaration of Helsinki.

### Reagents preparation

Fibrinogen (Sigma Aldrich, F4883, USA) 1 mg/ml in PBS was prepared as described in Shepeliuk et al^9^. Calcium-specific dye Calbryte-590^AM^ (AAT Bioquest, 20702, USA) was dissolved at 10 mM in DMSO, aliquoted, flash frozen in liquid nitrogen and stored at -80°C. On the day of experiment, one aliquot of Calbryte-590^AM^ was thawed at room temperature, diluted to 100 μM in Buffer A (150 mM NaCl, 2.7 mM KCl, 1 mM MgCl_2_, 0.4 mM NaH_2_PO_4_, 20 mM HEPES, 5 mM glucose, pH 7.4) and stored on ice in the dark until using in experiment. Annexin V-Alexa Fluor^647^ (BioLegend, 640912, CA) was stored at 4°C. Stock solutions of 10 mM mepacrine (Sigma Aldrich, Q3251, USA), thrombin at 20 NIH units (Sigma-Aldrich, T8885, USA) were prepared in milliQ water, aliquoted, flash frozen in liquid nitrogen, and stored at -80°C. On the day of the experiment, the relevant aliquots were thawed at room temperature, spun briefly and kept on ice until using in experiment but no longer than 1 h.

### Microfluidic platelet assay

Single-cell platelet visualization was carried out as in^9^. Briefly, plasma-treated coverslips were assembled with custom-made polydimethylsiloxane microfluidic chambers and functionalized by incubation with 1 mg/ml fibrinogen. Citrated whole blood was incubated with 1 μM mepacrine to label dense granules and Calbryte-590^AM^ to visualize intracellular calcium ions, and then flowed into a microfluidic channel at a wall shear rate of 1,000 s^−1^. Sparsely adhered platelets were activated by the addition of 0.1 U/ml thrombin in calcium-containing buffer and monitored in real time using Differential Interference Contrast (DIC) and fluorescence microscopy. Mepacrine and Calbryte-590^AM^ were excited by rapidly switching between 488 nm and 561 nm lasers (100 mW Sapphire lasers by Coherent, USA). The exposure time for each laser was 50 ms at 2 frames per second.

### Thrombodynamics assay

A portion of whole blood was centrifuged at 1600 × g for 15 min to obtain platelet-poor plasma (PPP). PPP was further centrifuged at 1,600 × g for 20 min to obtain platelet free plasma (PFP), which was used for a Thrombodynamics assay. The assay was performed using a Thrombodynamics Analyzer and reagent kit (HemaCore, LLC, Russia) as described previously^35^. This method measures spatial fibrin clot growth in a thin plasma layer initiated by contact with a surface bearing immobilized tissue factor. Clot formation was recorded by time-lapse imaging, and a fibrin growth curve was generated for each sample. Standard parameters included the steady clot growth rate, clot size and clot density at 30 min and the spontaneous clotting time (Tsp), defined as the time required for spontaneous clots to occupy 5% of the cuvette area.

### Experimental data analysis

Images were analyzed with ImageJ (National Institute of Health, USA). Data was analyzed using Origin (OriginLab Corporation, USA), Prizm (GraphPad Software, USA) and Matlab R2014a (MathWorks, USA) programs. Whenever possible, data are presented for N independent trials, i.e., experiments carried out in separate channels of a flow chamber using blood from the same donor/patient sample, to assess experimental reproducibility. In figure legends, lowercase n denotes the total number of analyzed platelets across all trials for the indicated conditions. Unless stated otherwise, values obtained in independent trials are reported as mean ± SD. For group-level comparisons, each patient or donor was treated as one biological replicate when applicable. Statistical significance was assessed using the Mann–Whitney U test; ns, not significant.

#### Quantification of the platelet adhesion and spreading

DIC images were analyzed to quantify platelet adhesion and morphology. For each experiment, ≥3 non-overlapping fields of view were selected before activation, and platelets within each field were counted. Adhesion density, expressed as cells per µm^2^, was calculated as the cell count divided by the field area (70 × 70 µm). Platelet morphology was scored by visual inspection: cells with a round-like shape, with or without filopodia, were classified as unspread, whereas cells with lamellipodia were classified as well-spread.

#### Quantification of the DG content in platelets

To determine DG content, approximately 10-30 platelets were selected from each field. Cells were selected using a semi-automatics approach (Supplementary Note 1). Briefly, regions of interest (ROIs) corresponding to individual platelets were identified using Trainable Weka Segmentation applied to the calcium channel (Calbryte-590^AM^) to generate binary masks. These ROIs were then applied to the mepacrine channel (488 nm) after background subtraction using a Gaussian blur (σ = 100). Total DG fluorescence per platelet was quantified as the integrated density within each ROI. ROIs with ambiguous morphology, including overlapping cells, edge artifacts, moving cells, or cells out of focus, were excluded based on DIC reference images.

#### Quantitative analysis of irregular calcium oscillations in individual platelets

Intracellular calcium dynamics were quantified from Calbryte-590^AM^ fluorescence time series, as described in Shepeliuk et al.^9^ Briefly, Calbryte-590^AM^ intensity was measured over time within a circular region inside each platelet. Calcium spikes were detected as local maxima above a manually selected threshold using a custom MATLAB script, which is available at https://www.med.upenn.edu/grishchuklab/protocols-software.html. Spike timing was extracted for each cell to calculate calcium spike frequency.

### Data availability

A Data Source file containing the quantitative data underlying all graphs is provided as the Supplementary Material. Due to their large size, raw imaging datasets (time-lapse microscopy sequences) are not included but are available from the corresponding author upon reasonable request.

## Results

### Quantitative fluorescence analysis resolves graded DG deficiency in pediatric patients

To evaluate DG abundance in pediatric patients diagnosed with DGD, blood samples from six patients were analyzed by electron microscopy (EM) using established clinical facilities. In five patients, the average number of DGs per platelet was reduced, falling below or close to the normal range of 4–6 DGs reported by this facility (Figure 2A). However, interpretation of the results was not always straightforward. Patient #2 was classified as normal because of the large SD in DG number, whereas patient #3, which exhibited a similar average DG count but a slightly smaller SD, was classified as DG-deficient. This discrepancy raised concerns about whether conventional summary statistics adequately capture DG content distributions.

**Figure 2.**
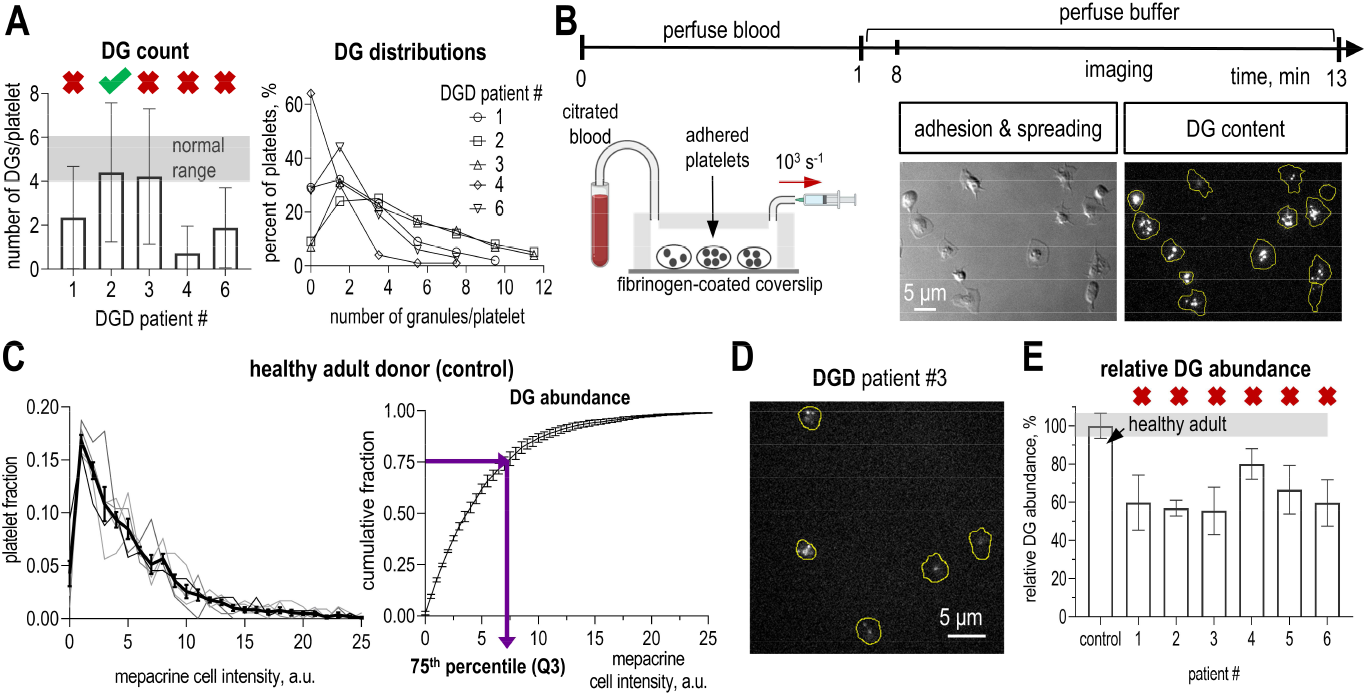
Single-platelet DG quantification in pediatric DGD patients. A. Clinical electron microscopy assessment of DGD patients. Left, DG counts per platelet reported by facility A. Bars show mean ±SD; the gray shaded area indicates the normal range established by the facility. Red crosses indicate patients classified as DG-deficient by the facility and green check marks indicate normal DG count. Right, DG-count distributions reported for individual DGD patients. B. Experimental workflow for functional platelet profiling. Citrated whole blood was perfused over fibrinogen-coated coverslips at 1,000 s^−1^ to immobilize platelets. After perfusion, platelets were washed with buffer, and adherent platelets were imaged to quantify adhesion and spreading, and DG content by mepacrine fluorescence. C. Single-platelet mepacrine fluorescence analysis in healthy adult donors used as control. Left, distributions of mepacrine fluorescence intensity in individual donors, reflecting platelet DG content. Thin gray curves show individual donors; the black curve shows the average distribution. Right, cumulative distribution of mepacrine fluorescence intensity. The 75^th^ percentile was used as a single-value metric of DG abundance. Total analyzed platelets: n = 1,490 from N = 6 healthy adult donors. D. Representative mepacrine fluorescence image of adherent platelets from DGD patient #3. Yellow contour indicates cell borders determined in corresponding DIC image. E. Relative DG abundance in DGD patients measured by single-platelet mepacrine fluorescence. The first bar shows the healthy adult reference set; subsequent bars show individual DGD patients analyzed in the same cohort as in panel A. Values are expressed as percent of the healthy adult control value, with n =30– 100 platelets analyzed for each patient. Uncertainty in this estimate was calculated by bootstrap resampling of individual patient platelet fluorescence values with replacement; error bars indicate bootstrap SD. Red crosses indicate patients classified as DG-deficient.

Inspection of the underlying EM data revealed that DG counts were not symmetrically distributed. Instead, all patient samples showed skewed distributions characterized by a reduced representation of platelets containing substantially more DGs than the population average (Figure 2A). As a result, mean and SD values alone provide only a limited description of DG abundance and may obscure biologically relevant differences between samples.

To characterize DG distributions in greater detail, we applied a quantitative fluorescence microscopy approach capable of measuring DG content in individual platelets while providing substantially larger sample sizes than are typically practical by EM. We used our previously developed microfluidic assay, in which citrated whole blood is perfused over fibrinogen-coated coverslips at 1,000 s^−1^, resulting in gentle immobilization of platelets without centrifugation or chemical fixation^9^ (Figure 2B; see Materials and Methods for details). DG content in adherent platelets was quantified from total cellular mepacrine fluorescence, an established marker of DG abundance^36,37^. Using a semi-automated image-analysis pipeline developed specifically for this application (Figure S1A-E; see Materials and Methods for details), we quantified both population-level features and platelet-to-platelet variability.

Analysis of healthy adult controls revealed that single-platelet mepacrine fluorescence follows a strongly non-Gaussian distribution, including a distinct subpopulation of DG-rich platelets (Figure 2C, total analyzed platelets n = 1,490 from N = 6 donors). This observation suggests that average fluorescence alone does not adequately capture the structure of the distribution. We therefore reconstructed cumulative fluorescence distributions and defined DG abundance as the 75th percentile of single-cell mepacrine fluorescence intensity (Figure 2C). This metric reflects the fluorescence level encompassing 75% of the platelet population while reducing sensitivity to extreme values and capturing changes in the DG-rich portion of the distribution.

When applied to the same DGD patient samples analyzed by EM, fluorescence measurements revealed reduced DG content in every patient relative to healthy controls, with values ranging from 55% to 80% of the control level (Figure 2A,D,E). Importantly, the method resolved graded differences among patients rather than assigning samples to binary normal/deficient categories. An additional patient was evaluated by EM at a different clinical facility, where any DG number above 1.2 per platelet was considered normal (Figure S1F). Although this patient was classified as normal by EM, fluorescence analysis detected a significant reduction in DG content. Together, these findings indicate that quantitative fluorescence measurements provide a sensitive framework for evaluating DG abundance, revealing deficiencies and patient-to-patient differences that are not fully captured by conventional clinical EM classification.

### Reduced DG abundance extends across multiple pediatric platelet disorders

To further evaluate the performance of the assay, we analyzed reference samples from patients with Gray platelet syndrome and MYH9 disorder, in which reduced and elevated DG content, respectively, were expected. Total mepacrine fluorescence was quantified and cumulative distributions were reconstructed. Both samples differed markedly from healthy controls. The Gray platelet syndrome sample exhibited a substantially steeper cumulative distribution, whereas the MYH9 sample showed a pronounced downward shift (Figure 3A,B, Figure S2). These findings confirm that the assay is sensitive across a clinically relevant range of DG abundance.

**Figure 3.**
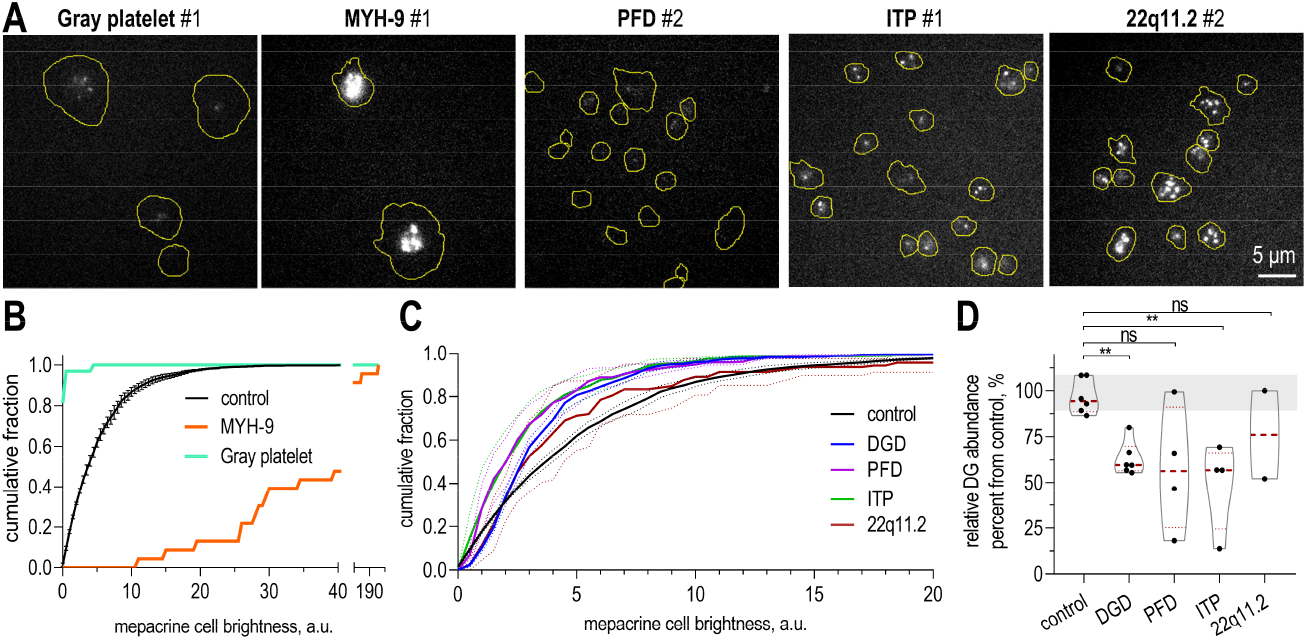
Diverse DG content across pediatric bleeding disorders. A. Representative mepacrine fluorescence images of adherent platelets from pediatric patients with different platelet disorders. Yellow contours indicate cell contours. B. Examples of pronounced DG depletion and elevated DG content in individual patients. Cumulative distributions of single-platelet mepacrine fluorescence intensity are shown for healthy adult controls, a Gray platelet syndrome patient with severely reduced DG content, and a MYH-9 patient with increased DG content. N and n. C. Cumulative distributions of single-platelet mepacrine fluorescence intensity in healthy adult controls and color-coded pediatric diagnostic groups. Solid lines represent average group distributions of platelet mepacrine intensity; dotted lines indicate SEM. D. Relative DG abundance in the same patients as in panel C. Each dot represents one donor or patient, based on 20–100 analyzed platelets per patient. The gray band indicates the healthy adult control range; red dashed lines indicate group means. Statistical comparisons were performed using Mann–Whitney tests; ns, not significant; **p < 0.01.

We next examined whether altered DG content was restricted to patients diagnosed with DGD or was also present in other pediatric platelet disorders in which DG abundance has not been systematically assessed using high-resolution single-cell measurements. Platelets from patients with platelet function defects (PFD), immune thrombocytopenia (ITP), and 22q11.2 deletion syndrome were immobilized and analyzed using the same approach (2–9 patients per group). Clinical characteristics of the patient groups are summarized in Supplementary Table 1, but briefly, patients ranged from 1 to 22 years of age and showed broad variation in platelet counts, while platelet volumes were mostly within the normal range when available. Most patients exhibited only mild bleeding manifestations despite their underlying diagnoses, with bleeding scores of 0–1 across the groups. Available clinical platelet testing showed heterogeneous abnormalities, including abnormal aggregation and/or impaired secretion responses. Single-cell mepacrine imaging of these platelets revealed a broad range of DG abundance among individual platelets (Figure 3A), while cumulative population distributions showed reduced DG content relative to healthy adult controls in both PFD and ITP cohorts and a more modest shift in 22q11.2 deletion syndrome (Figure 3C).

To quantify inter-patient variability, DG abundance was expressed relative to healthy control platelets, which were assigned an average value of 100%. Healthy donors ranged from approximately 87.4% to 107.4%, establishing a reference interval for normal variation (Figure 3D). Using this metric, average DG abundance was reduced in all examined patient groups, typically to 55–65% of control values, comparable to the reductions observed in the DGD cohort. However, substantial variability was present within each diagnostic category (Figure 3D, Figure S3). Some PFD patients exhibited only 18–66% of normal DG abundance, whereas one patient remained within or near the healthy reference range. All analyzed ITP patients showed reduced DG abundance, including one patient with only 14% of normal content, markedly lower than any patient in the DGD cohort. Among the two patients with 22q11.2 deletion syndrome, one exhibited approximately 50% reduced DG abundance, whereas the other remained within the healthy adult range.

Together, these results demonstrate that reduced DG content is not confined to patients diagnosed with DGD. Instead, diminished DG abundance occurs across multiple pediatric platelet disorders and spans a broad continuum of severity, from near-normal levels to deficiencies exceeding those observed in some DGD patients. This variability is not fully explained by current diagnostic categories and suggests that DG-related phenotypes may cut across clinically distinct platelet disorders.

### Platelet adhesion and spreading vary independently of diagnostic category

DG abundance varied substantially among patients with different platelet disorders. We therefore asked whether other platelet functional properties exhibited similar variability and whether these features aligned with diagnostic category. As a functional readout, we examined platelet adhesion and spreading on fibrinogen-coated coverslips^9^. After perfusion of whole blood for 1 min, blood was replaced with buffer and flow chambers were imaged after an additional 7 min using differential interference contrast microscopy (Figure 4A). The number of adherent platelets and the fraction of platelets exhibiting prominent lamellipodia were quantified.

**Figure 4.**
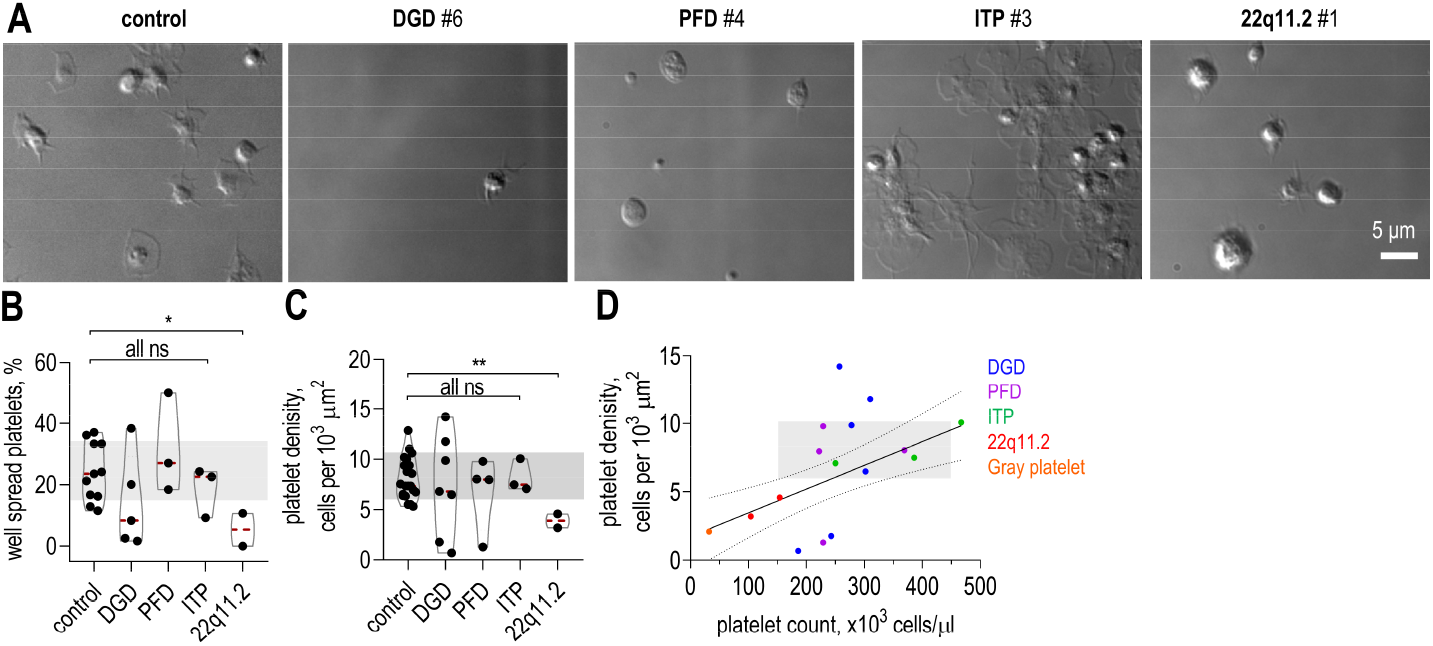
Diverse platelet adhesion and spreading across pediatric bleeding disorders. A. Representative DIC images of platelets adhered to fibrinogen-coated coverslips under flow in healthy adult control blood and pediatric patient samples. B. Platelet spreading across healthy adult controls and pediatric diagnostic groups. The percent of well-spread platelets was quantified based on DIC morphology; cells with visible lamellipodia were counted as well spread. Here and in panels C,D, each dot represents one healthy donor or patient; the gray band indicates the healthy adult control range. Statistical comparisons were performed relative to healthy controls using Mann–Whitney test; ns, not significant; *p < 0.05. C. Platelet adhesion quantified as the surface density of adherent cells across healthy controls and pediatric diagnostic groups. See panel B for other details. Statistical comparisons were performed relative to healthy controls using Mann–Whitney test; ns, not significant; **p < 0.01. D. Relationship between platelet count and platelet adhesion density across pediatric patient samples. Each dot represents one patient and is color-coded by diagnostic group. The gray shaded area indicates the healthy adult reference ranges for both platelet count (150–450 ×10^3^ platelets/μl) and platelet adhesion density. The solid line shows the linear regression fit, with dotted lines indicating the 99% confidence interval. Pearson correlation coefficient is 0.97. Several samples fall outside the confidence interval, indicating patient-specific deviations from the overall relationship between platelet count and adhesion density.

The proportion of spread platelets with lamellipodia varied considerably even among healthy adult controls, ranging more than threefold from 11.5% to 37% (Figure 4B, Figure S4A). Similar variability was observed across most diagnostic groups. In contrast, both patients with 22q11.2 deletion syndrome exhibited comparably reduced spreading. DGD patients also tended to show lower spreading than controls. However, given the limited cohort sizes, average differences among DGD, PFD, and ITP groups were not statistically significant, and it was not possible to determine whether variability differed systematically among diagnostic categories. Instead, spreading behavior appeared to be largely patient-specific and not explained by diagnosis alone (Figure 4B).

We next examined platelet adhesion. The surface density of adherent platelets varied most prominently within the DGD cohort, ranging from only 1 to 15 cells/µm^2^, whereas healthy controls exhibited 8.1 ± 2.1 cells/µm^2^ (Figure 4C, Figure S4B). ITP samples generally showed preserved adhesion, although one patient (#3) displayed an unusually large number of platelet aggregates (Figure 4A,C). In contrast, adhesion appeared reduced in both patients with 22q11.2 deletion syndrome. Importantly, adhesion and spreading phenotypes did not fully overlap. Some samples exhibited reduced spreading despite relatively preserved adhesion, indicating that platelet attachment and spreading represent related but partially independent functional properties.

Variation in platelet adhesion density can arise simply from differences in circulating platelet number. To distinguish between these possibilities, we plotted the density of adherent platelets against platelet counts measured in the same blood samples (Figure 4D). Interestingly, samples from patients with 22q11.2 deletion syndrome, which exhibited the lowest adhesion densities, as well as most ITP samples, fell close to a common linear relationship between platelet count and surface density. This finding suggests that adhesion propensity in these groups can largely be explained by numbers of circulating platelets rather than altered adhesive behavior per platelet. In contrast, while some DGD and PFD samples also mapped within the expected range, several deviated substantially from this relationship. Certain patients exhibited more adherent platelets than predicted by platelet count alone, whereas others exhibited fewer, suggesting increased or decreased adhesive propensity independent of platelet abundance (Figure 4D).

Together, these findings indicate that platelet adhesion and spreading constitute an additional functional axis that varies across pediatric platelet disorders. While some differences can be attributed to platelet count, others likely reflect intrinsic alterations in platelet behavior that are not captured by conventional diagnostic classification.

### Calcium signaling reveals heterogeneous platelet activation states across pediatric disorders

DG abundance, spreading, and adhesion revealed substantial patient-to-patient variability. However, these measurements characterize platelets in the absence of exogenous stimulation. We therefore asked whether additional differences emerge during agonist-induced activation. As a dynamic readout of platelet activation, we examined intracellular calcium signaling following thrombin stimulation. Thrombin is known to induce irregular calcium oscillations whose frequency reflects the degree of platelet activation^9^.

Platelets in whole-blood samples were labeled with Calbryte-590^AM^ and perfused through microfluidic chambers^9^. After adhesion to fibrinogen-coated coverslips and washing with buffer, adherent platelets were stimulated with 0.1 U/mL thrombin and monitored by fluorescence microscopy (Figure 5A). Before thrombin addition, healthy adult platelets exhibited infrequent and irregular calcium spikes^9^. Following stimulation, the stochastic character of calcium signaling was preserved, but spike frequency increased markedly (Figure 5B). We therefore quantified average calcium spike frequency over the subsequent 9 min interval, as described previously^9^. Measurements from healthy adult donors followed an approximately normal distribution with an average frequency of 13.1 ± 2.3 spikes/min, establishing a physiological reference range for thrombin-induced calcium activity (Figure 5C).

**Figure 5.**
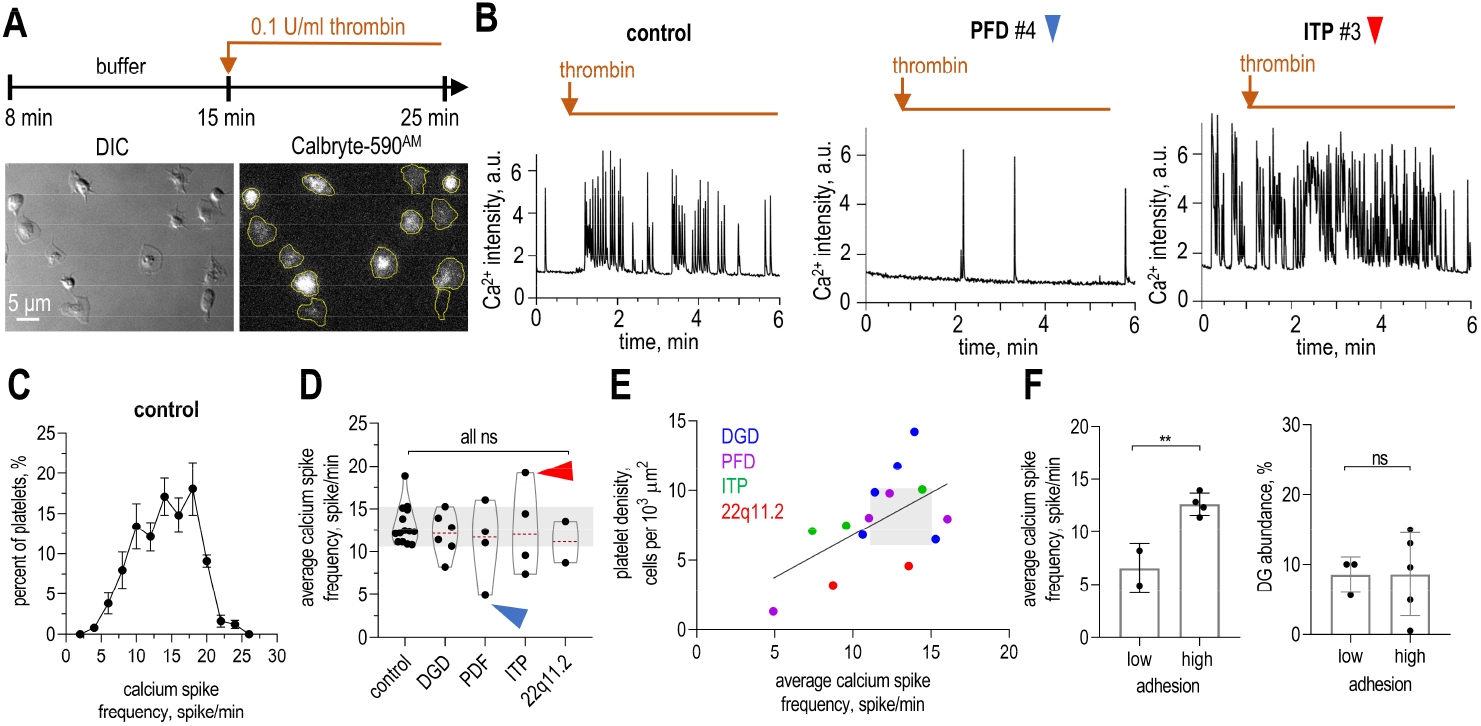
Diverse calcium signaling across pediatric bleeding disorders. A. Experimental workflow of platelet activation and representative images for single-platelet calcium imaging. Continuing the timeline shown in Figure 2B, adherent platelets were stimulated with 0.1 U/mL thrombin. DIC and Calbryte-590^AM^ fluorescence images show the same field of immobilized platelets with yellow contour indicating cell contour. B. Representative Ca^2^+ fluorescence trace following addition of thrombin (0.1 U/mL) in healthy adults and pediatric patient samples. Orange lines indicate the period of thrombin exposure. C. Distribution of calcium spike frequencies in healthy adult control platelets. Data are shown as mean ± SEM for N=5 and n=241. D. Calcium spike frequency across healthy adult controls and pediatric diagnostic groups. Each dot represents one healthy donor or patient; the gray band indicates the healthy adult control range; red dashed lines indicate group means based on 20–50 analyzed platelets per patient. Statistical comparisons were performed relative to healthy controls using Mann–Whitney tests; ns, not significant. E. Relationship between calcium spike frequency and platelet adhesion density across pediatric patient samples. Each dot represents one patient and is color-coded by diagnostic group. The solid line shows the linear regression fit. The gray shaded area indicates the healthy adult reference ranges for both calcium spike frequency and platelet adhesion density. Pearson correlation coefficient is 0.63. F. Relationship between platelet adhesion and calcium signaling or DG abundance. Left, calcium spike frequency; right, relative DG abundance in low- and high-adhesion patient samples. Low- and high-adhesion groups were defined based on deviations from the platelet count–adhesion relationship shown in Figure 4E, using the confidence interval as the cutoff. Bars show mean ± SEM; each dot represents one patient. Statistical comparisons were performed using Mann–Whitney tests; ns, not significant; **p < 0.01.

We next applied the same analysis to platelets from pediatric patients (Supplementary Table 1). Most samples exhibited calcium spike frequencies within the healthy adult range (Figure 5D, Figure S4C). However, several notable deviations were observed. PFD patient #4 showed near-complete suppression of calcium spiking, whereas ITP patient #3 displayed markedly elevated spike frequency (Figure 5B). Another ITP patient exhibited below-normal activity (Figure 5D). Thus, both hypo-reactive and hyper-reactive platelet activation states were detected across the cohort, including within the same diagnostic category.

Because platelet calcium signaling can be closely linked to adhesive interactions^38^, we next compared calcium spike frequency with platelet adhesion measured in the same chambers. Across all pediatric samples, calcium spike frequency showed a positive relationship with platelet adhesion density (Figure 5E, Figure S5), suggesting that these parameters reflect related aspects of platelet activation state. To further examine this relationship, we focused on samples in which adhesion was noticeably increased or decreased relative to the value expected from platelet count alone. Consistent with the overall trend, platelets with elevated adhesive propensity exhibited nearly twofold higher calcium spike frequencies than platelets with reduced adhesion (Figure 5F, Figure S5). In contrast, DG abundance did not differ between these groups despite the pronounced differences in adhesion and calcium activity. This observation is consistent with the expectation that DG content in resting platelets primarily reflects granule abundance rather than activation state.

Together, these results indicate that different platelet phenotypes capture distinct aspects of platelet biology. While DG abundance, adhesion/spreading behavior, and calcium signaling can each be altered in pediatric platelet disorders, these features do not vary in a coordinated manner across patients. Instead, individual patients exhibit distinct combinations of abnormalities, revealing functional subphenotypes that are not readily captured by conventional diagnostic classification or population-averaged measurements.

### Enhanced plasma coagulation despite platelet functional abnormalities

Many patients exhibited altered platelet functional readouts, including reduced DG abundance, impaired spreading or adhesion, and abnormal calcium signaling, yet most had only minimal bleeding manifestations (Supplementary Table 1). This observation raised the possibility that plasma coagulation might compensate, at least in part, for impaired platelet function. To address this question, we analyzed platelet-free plasma from the same pediatric cohorts using the Thrombodynamics assay, which quantifies fibrin clot propagation from a localized tissue factor-bearing surface. Unlike conventional global coagulation tests, Thrombodynamics captures the spatial nature of coagulation by measuring clot growth as a propagating process originating from a defined activation site rather than from a uniformly distributed activator^35^. Briefly, CTI-treated, recalcified citrated plasma was placed into a cuvette and brought into contact with a tissue factor-coated activator insert, mimicking a localized site of vascular injury (Figure 6A,B). Fibrin clot formation initiated at the tissue factor surface and propagated through the plasma over time, enabling quantification of clot growth kinetics (Figure 6C). In addition, deregulated coagulation can be detected through spontaneous clot formation away from the activation surface. Because of its high reproducibility and low sample-volume requirements, this assay is particularly well suited for pediatric studies^35,39–47^.

**Figure 6.**
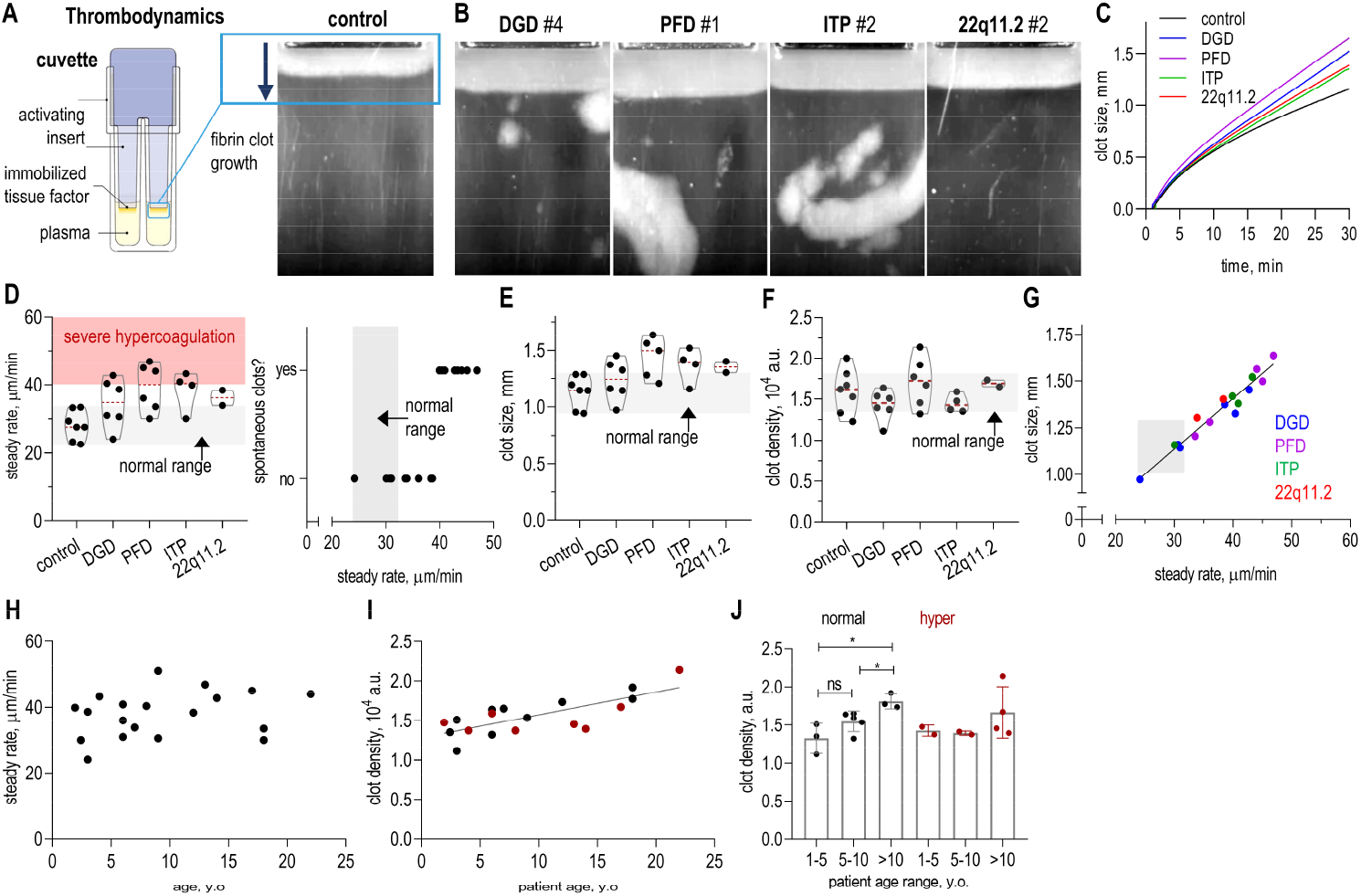
Altered fibrin clot growth dynamics in pediatric bleeding disorders. A. Schematic of the Thrombodynamics assay. Platelet-free plasma was placed in a cuvette, and fibrin clot growth was initiated from immobilized tissue factor at the activating insert. Clot propagation was monitored over time to quantify growth rate, clot size, and clot density. A representative clot from a healthy adult control is shown at 30 min; the tissue factor surface is located at the upper edge of the image. Vertical lines are the scratches on the surface of the plastic cuvette. B. Representative fibrin clot images at 30 min in pediatric patient plasma. Images illustrate differences in clot propagation in comparison to control clot in panel A and the presence or absence of spontaneous clotting away from the activating surface. C. Representative clot growth curves in healthy adult control and color-coded pediatric diagnostic groups. Clot size was measured over time from the tissue factor surface. D. Steady-state clot growth rate and spontaneous clotting across healthy adult controls and pediatric diagnostic groups. Left, steady-state clot growth rate; each dot represents one donor or patient, and red dashed lines indicate group means. The gray shaded region indicates the normal reference range, and the red shaded region indicates the hypercoagulable range associated with spontaneous clotting. Right, presence or absence of spontaneous clotting plotted against steady-state clot growth rate. E. Clot size at 30 mins. See panel D for details. F. Clot density at 30 mins. See panel D for details. G. Relationship between clot size and steady-state clot growth rate across pediatric patient samples. Each dot represents one patient and is color-coded by diagnostic group. The gray shaded area indicates the healthy adult reference ranges for both steady rate and clot size. The solid line shows the linear regression fit. Pearson correlation coefficient is 0.97. H. Relationship between steady-state clot growth rate and patient age. Each dot represents one pediatric patient. Pearson correlation coefficient is 0.26. I. Relationship between clot density and patient age. Each dot represents average for one pediatric patient. Black points indicate patients within the normal coagulation range, and red points indicate patients classified as hypercoagulable with spontaneous clotting. Pearson correlation coefficient is 0.75. J. Clot density grouped by age range and coagulation status. Bars show mean ± SEM. Statistical comparisons were performed relative to healthy controls using Mann–Whitney test; ns, not significant; *p < 0.05. See panel I for details.

Plasma coagulation measurements were performed on the same blood samples used for platelet functional analysis, minimizing variability associated with blood collection and sample processing. We quantified steady rate of clot growth, clot size throughout propagation, final clot size, and optical clot density 30 min after initiation of coagulation. In healthy adult controls, clot growth rate exhibited a relatively narrow distribution centered around 28.2 ± 4.3 µm/min (Figure 6D). Strikingly, plasma coagulation was frequently elevated in pediatric patients. Across all diagnostic groups, average clot growth rates exceeded those observed in healthy controls (Figure 6B–D). Several individual patients with DGD, PFD, and ITP exhibited clot growth rates above 40 µm/min, a threshold considered indicative of hypercoagulability. Final clot size was likewise frequently increased (Figure 6E). In contrast, clot density exhibited more variable behavior, with most samples remaining within or near the healthy adult reference range (Figure 6F). Across all patient groups, final clot size correlated strongly with clot growth rate (Figure 6G), indicating that propagation velocity remains a major determinant of clot size. This relationship appears preserved in these pediatric disorders despite substantial variation in coagulation activity.

As clot growth rate from the activation surface increased, spontaneous fibrin clot formation away from the activator became increasingly apparent (Figure 6D, Figure S6), providing independent evidence of hypercoagulable plasma behavior. Importantly, both normal and hypercoagulable samples were observed within the same diagnostic categories. Despite the frequent occurrence of hypercoagulation, we found no obvious relationship between coagulation parameters and any of the platelet phenotypes measured in the same samples, including DG abundance, adhesion, spreading, or calcium signaling (Figure S7). Although the cohort size limits formal correlation analysis, these observations suggest that plasma coagulation represents an additional source of functional variability that is not directly captured by platelet-centered measurements alone.

Additional complexity emerged when coagulation parameters were analyzed as a function of patient age. Clot density showed a positive age-associated trend, whereas steady-state clot growth rate exhibited no clear age dependence (Figure 6H,I). Previous studies have reported age-dependent changes in fibrinogen concentration and fibrin network architecture^48,49^, and our measurements are consistent with this general pattern. Notably, the age-dependent increase in clot density was most evident among samples with otherwise normal coagulation parameters (Figure 6J). In contrast, among hypercoagulable samples, this trend was less apparent, suggesting that disease-associated alterations in coagulation may partially obscure underlying developmental effects. Together, these findings reveal an unexpected dissociation between platelet functional abnormalities and plasma coagulation behavior in pediatric platelet disorders. Impaired platelet function frequently coexisted with enhanced plasma coagulation, including hypercoagulable phenotypes that were not readily explained by diagnostic category or by the platelet parameters measured here.

## Discussion

Pediatric platelet disorders encompass diverse disease mechanisms, yet their functional phenotypes remain incompletely understood. In this pilot study, we combined quantitative platelet and coagulation measurements across several pediatric diagnostic groups and identified substantial variability in multiple functional readouts (Figure S8). While the limited cohort size and heterogeneous patient population preclude definitive disease-specific conclusions, the study revealed unexpected phenotypes, exposed important gaps in current functional characterization, and highlighted opportunities for more comprehensive future investigations.

A notable outcome of this study was the development and validation of a quantitative single-cell assay for DG abundance. Unlike conventional EM-based analysis, the fluorescence approach allowed hundreds of individual platelets from each patient sample to be analyzed under minimally perturbing conditions^9^, providing sufficient statistical power to characterize distributional features of DG content. This analysis revealed that DG abundance in healthy platelets follows a strongly non-Gaussian distribution, making conventional summary statistics such as mean and SD insufficient to describe the population. By introducing a percentile-based metric of DG abundance and validating it using DGD, Gray platelet syndrome, and MYH9 reference samples, we established a sensitive framework for quantitative assessment of platelet granule content. Beyond its application in the present study, this approach may facilitate future investigations of platelet granule biology in both pediatric and adult cohorts.

Application of this assay revealed an unexpected finding. As anticipated, DGD patients exhibited uniformly reduced DG abundance. However, reduced DG content was also observed in several patients with PFD and ITP, disorders in which granule deficiency is not typically considered a defining feature. These observations suggest that diminished DG abundance may represent a broader characteristic of pediatric platelet dysfunction than currently appreciated. Importantly, the ability to detect these differences likely reflects both the improved quantitative resolution of the assay and the larger numbers of platelets analyzed compared with conventional EM-based approaches.

The most striking observation, however, was the extent of functional diversity observed within diagnostic categories. While some disease groups exhibited modest shifts in average behavior, group means often obscured substantial patient-to-patient variability. Individual patients displayed distinct combinations of altered DG abundance, adhesion, spreading, and calcium signaling, indicating that these measurements capture partially independent aspects of platelet biology. For example, the two patients with 22q11.2 deletion syndrome both exhibited reduced adhesion, consistent with altered GPIb-dependent platelet function, yet differed markedly in other platelet phenotypes. Similar divergence was observed in PFD and ITP cohorts, where patients sharing the same clinical diagnosis displayed markedly different activation and signaling profiles. Collectively, these findings suggest that diagnosis alone does not reliably predict platelet functional state and that multidimensional phenotyping may provide a more informative description of platelet dysfunction than any individual assay.

Future studies should expand these observations in larger patient cohorts and through longitudinal sampling of individual patients. Such studies should integrate clinical history, treatment status, and additional platelet phenotypes, particularly direct measurements of secretion dynamics. Granule abundance and granule release represent mechanistically distinct processes and may contribute differently to overall hemostatic function. Time-resolved measurements of secretion therefore represent an important next step toward understanding how diverse platelet phenotypes contribute to clinical manifestations in pediatric bleeding disorders.

The most unexpected finding of this study emerged from plasma clotting measurements. Using the Thrombodynamics assay, we observed frequent hypercoagulable phenotypes characterized by accelerated clot growth and, in some patients, spontaneous fibrin formation despite clinical diagnoses associated primarily with platelet-related bleeding disorders. This observation was particularly striking because several patients exhibited impaired platelet phenotypes while simultaneously demonstrating enhanced plasma clotting potential. Although the prevalence of hypercoagulation varied among diagnostic groups and cohort sizes were limited, these findings suggest that enhanced plasma clotting may be more common in pediatric platelet disorders than currently appreciated.

The ability to detect these phenotypes highlights the value of the Thrombodynamics approach. Unlike conventional coagulation tests, the assay reproduces spatial propagation of clotting from a localized tissue factor source and simultaneously reports multiple parameters of clot formation, including propagation rate, clot size, clot density, and spontaneous fibrin generation. Importantly for pediatric studies, these measurements can be performed using small plasma volumes obtained from the same blood samples used for platelet functional analysis, enabling integrated characterization of both cellular and plasma components of hemostasis.

The present study was not designed to determine the origin of the hypercoagulable phenotype. Potential contributors include compensatory adaptations to platelet dysfunction, disease-associated inflammatory signaling, developmental effects, treatment history, or other patient-specific factors. We did not observe a clear relationship between plasma clotting parameters and the platelet phenotypes measured here, although interpretation is limited by cohort size, age heterogeneity, multiple diagnostic categories, and single-time-point sampling. Nevertheless, these findings demonstrate that overall hemostatic potential cannot be inferred from platelet abnormalities alone and emphasize the importance of evaluating platelet and plasma pathways together, even in disorders traditionally viewed as primarily platelet-mediated.

More broadly, our results highlight the need for increasingly physiological functional assays capable of integrating platelet activity and plasma coagulation within a common experimental framework. Future studies should incorporate real-time measurements of whole-blood clot formation under vessel-mimicking flow conditions to determine how the diverse platelet and plasma phenotypes identified here combine to shape overall hemostatic function.

Another important observation emerged from the analysis of age-dependent effects. We found that fibrin clot density increased with age, consistent with previous reports linking maturation of the hemostatic system to changes in fibrinogen levels and fibrin network structure^48,49^. Although the magnitude of this effect was generally smaller than many disease-associated changes observed in our cohort, its detection provides an important validation of the sensitivity of the Thrombodynamics assay. Interestingly, age-related trends were evident for clot density but not for other clotting parameters, suggesting that developmental influences may affect different aspects of coagulation to different degrees.

More broadly, these findings highlight a major challenge in the interpretation of pediatric hemostatic phenotypes. During infancy, substantial differences in coagulation factor levels and activities relative to adults are well established as part of developmental hemostasis^50,51^. Beyond infancy, however, quantitative reference frameworks remain surprisingly limited. Standard clinical assays often report broadly similar results across age groups^50,52–54^, yet more detailed studies have identified age-dependent differences in platelet granule content, receptor expression, activation responses, and other functional parameters^55–59^. Existing reports are relatively sparse and often difficult to compare because of differences in methodology and cohort composition. Consequently, interpretation of functional abnormalities in pediatric patients remains constrained by the absence of robust quantitative baselines established specifically for healthy children^3,60^. Like many previous studies, our work used healthy adults as controls. While this limitation is unlikely to affect the major conclusions of the present study, particularly comparisons among patient cohorts analyzed under identical conditions, future investigations will require age-matched pediatric reference populations. Establishing quantitative baselines for platelet phenotypes and plasma clotting parameters across childhood should become a priority for the field and will be essential for distinguishing pathological alterations from normal developmental variation.

In conclusion, this pilot study demonstrates the value of combining quantitative single-cell platelet phenotyping with spatially resolved measurements of plasma clotting. The approach revealed reduced dense granule abundance outside classical DGD, substantial functional diversity within established diagnostic categories, and unexpectedly frequent hypercoagulable plasma phenotypes in patients evaluated for platelet-related bleeding disorders. While the molecular origins and clinical significance of these findings remain to be determined, the study highlights important gaps in current understanding, identifies several areas in which diagnostic and experimental approaches can be improved, and provides a foundation for future investigations of pediatric hemostasis. More broadly, our results argue that platelet and plasma pathways should be studied together using quantitative, physiologically relevant assays capable of capturing the multidimensional nature of hemostatic function.

## Supporting information

Supplementary File with Figures, Tables, Notes

## Acknowledgements

We thank Grishchuk lab members for discussions and assistance. Research conducted in the Grishchuk laboratory was supported by internal research funds from the Perelman School of Medicine at the University of Pennsylvania. Masks were prepared at Singh Center for Nanotechnology (University of Pennsylvania, USA), which is supported by the NSF National Nanotechnology Coordinated Infrastructure Program under grant NNCI-2025608. Chamber holders were made at 3D printing service at Biotech Commons (University of Pennsylvania, USA).

## Authorship contributions

T.O.S performed experiments in vitro and analyzed data, P.K. collected pediatric samples and clinical data, E.M., E.H and F.A. developed protocols and software for semi-automatic analysis of DG content, T.O.S, M.P.L and E.L.G designed research and interpreted data, T.O.S. and E.L.G. wrote the manuscript with input from other co-authors.

