## Supplementary File with Figures, Tables, Notes for "Hidden Complexity of Pediatric Platelet Disorders: Functional Diversity and Unexpected Hypercoagulable Phenotypes"

**TABLE OF CONTENT**


1. Supplementary Tables
2. Supplementary Figures and Figure Legends
3. Supplementary Notes
4. Supplementary References

#### Supplementary Tables

##### Table 1. Clinical characteristics of pediatric patients included in the study.

| Patient ID | Assigned group | Age, y.o. | Sex | Platelet count, ×10³/μL | MPV, fL | Bleeding score | Clinical description | Clinical platelet testing |
|---|---|---|---|---|---|---|---|---|
| **DGD #1** | DGD | 6 | M | 257 | 10.0 | 0 | history of bleeding and family history of an inherited platelet disorder. | EM DG count = 2.19 Normal aggregation and secretion |
| **DGD #2** | DGD | 3 | F | 310 | 9.3 | 0 | family history of bleeding disorder, no significant past medical history. | EM DG count = 4.38 Normal aggregation and secretion |
| **DGD #3** | DGD | 5 | M | 243 | 9.3 | 1 | family history of bleeding disorder, Maternal history of platelet storage defect. | EM DG count = 4.22, reduced secretion to ADP only |
| **DGD #4** | DGD | 14 | F | 278 | 9.0 | 0 | delta granule storage pool disorder | EM DG count = 0.64, reduced secretion to ADP only |
| **DGD #5** | DGD | 9 | F | 302 | 10.5 | 0 | frequent nosebleeds and need platelet aggregation testing | EM DG count = 3.2 Normal aggregation and secretion |
| **DGD #6** | DGD | 8 | F | 186 | 9.9 | 0 | delta granule storage pool deficiency | EM DG count = 1.79 Normal aggregation and secretion |
| **PFD #1** | PFD | 13 | M | 222 | 11.6 | 1 | platelet function defect | normal aggregation and secretion |
| **PFD #2** | PFD | 17 | F | 229 | 10.7 | 0 | qualitative platelet function defect | abnormal secretion to ADP and collagen and abnormal agglutination in response to ristocetin |
| **PFD #3** | PFD | 22 | F | 306 | 10.5 | 0 | platelet function defect | NA |
| **PFD #4** | PFD | 18 | F | 229 | 10.1 | 0 | platelet function defect | reduced aggregation in response to collagen, AA, ristocetin, reduced secretion in response to ADP and collagen |
| **PFD #5** | PFD | 7 | F | 369 | 8.5 | 0 | abnormal platelet function test, Frequent nosebleeds | normal aggregation and secretion |
| **ITP #1** | ITP | 4 | F | 467 | 9.6 | 0 | immune thrombocytopenia | NA |
| **ITP #2** | ITP | 6 | M | 250 | 9.6 | 0 | immune thrombocytopenia, RAS associated lymphoproliferative disease | NA |
| **ITP #3** | ITP | 2.4 | F | 352 | 8.2 | 0 | immune thrombocytopenia | reduced aggregation and secretion in response to ADP only |
| **ITP #4** | ITP | 2 | F | 386 | 8.9 | 0 | immune thrombocytopenia | NA |
| **22q11.2 #1** | 22q11.2 deletion | 7 | M | 104 | NA | 0 | 22q11.2 deletion syndrome | NA |
| **22q11.2 #1** | 22q11.2 deletion | 12 | F | 154 | NA | 0 | 22q11.2 deletion syndrome | NA |
| **MYH-9 #1** | MYH-9 disorder | 9 | M | 15 | NA | 0 | familial macrothrombocytopenia (suspected MYH9 related) | EM DG count = 9.57 |
| **Gray platelet #1** | Gray platelet syndrome | 7 | NA | 32 | NA | 0 | macrothrombocytopenia with possible gray platelet syndrome | NA |

Abbreviations: ITP – immune thrombocytopenia, DGD – dense granule deficiency, PFD – platelet function defect, NA – not available.

### Supplementary Figures and Figure Legends

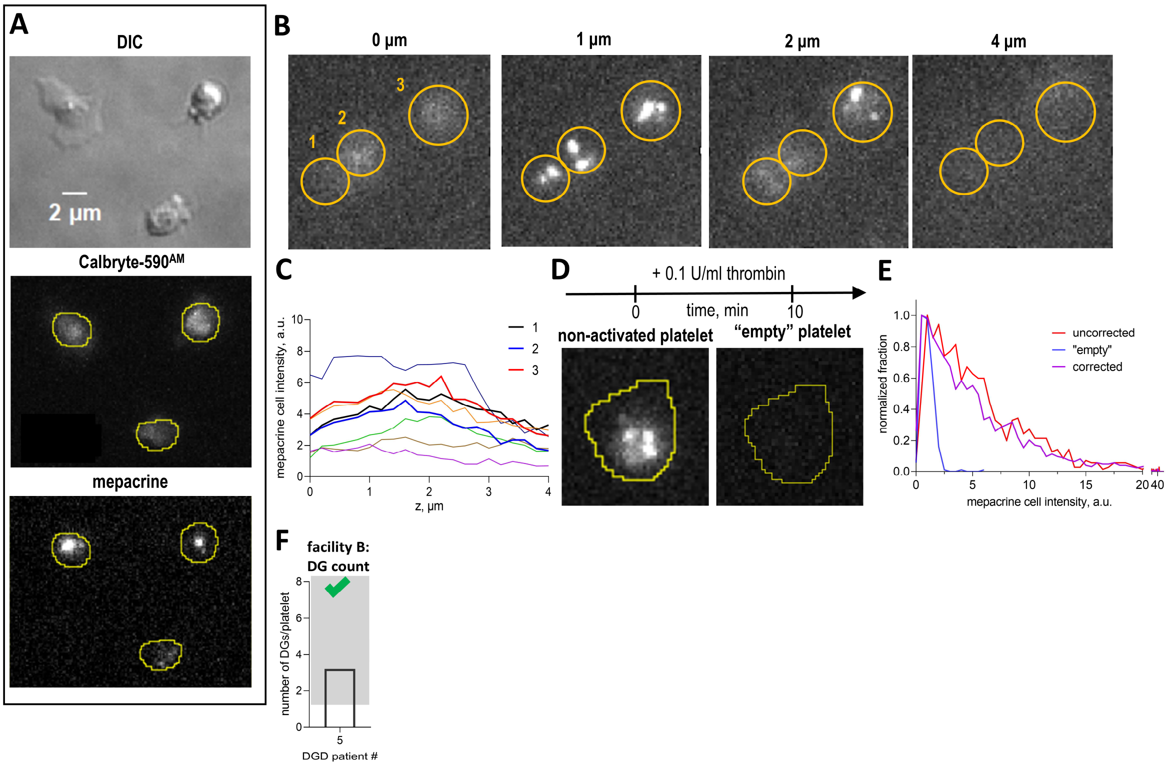


**Supplementary Figure 1. Semi-automated image analysis of mepacrine-based DG quantification.**

A. Representative images of immobilized platelets acquired by DIC microscopy, Calbryte-590$^{AM}$ fluorescence, and mepacrine fluorescence. Yellow contours indicate segmented platelet contours used for mepacrine fluorescence analysis.

B. Representative z-stack images of mepacrine labeled platelets. The same platelets are shown at different z-positions to illustrate how fluorescence intensity varies through the platelet volume. Yellow circles indicate platelet location.

C. Mepacrine fluorescence intensity across the z-stack for individual platelets. Each curve represents one segmented platelet, showing variation in total platelet fluorescence across z-position. Bold curves indicate platelets shown in panel B.

D. Example of mepacrine fluorescence before and after platelet activation. A non-activated platelet retains bright mepacrine-positive DG signal, whereas an activated platelet shows loss of mepacrine fluorescence and appears as an “empty” platelet used for analysis.

E. Distribution of mepacrine fluorescence intensity in healthy adult platelets before and after correction for mepacrine fluorescence in “empty” platelets. Red curve shows non-corrected mepacrine fluorescence of platelets (N=28, n=1,235), blue curve shows the fluorescence of “empty” platelets (N=17, n=245), and magenta curve shows the corrected distribution after exclusion of distribution of empty platelets (N=30, n=1490).

F. Clinical electron microscopy assessment of DGD patient. DG counts per platelet reported for patient #5 by another facility than in Figure 2A. The bar shows the mean; SD is not shown because it was not reported. The gray shaded area indicates the normal range established by the facility.

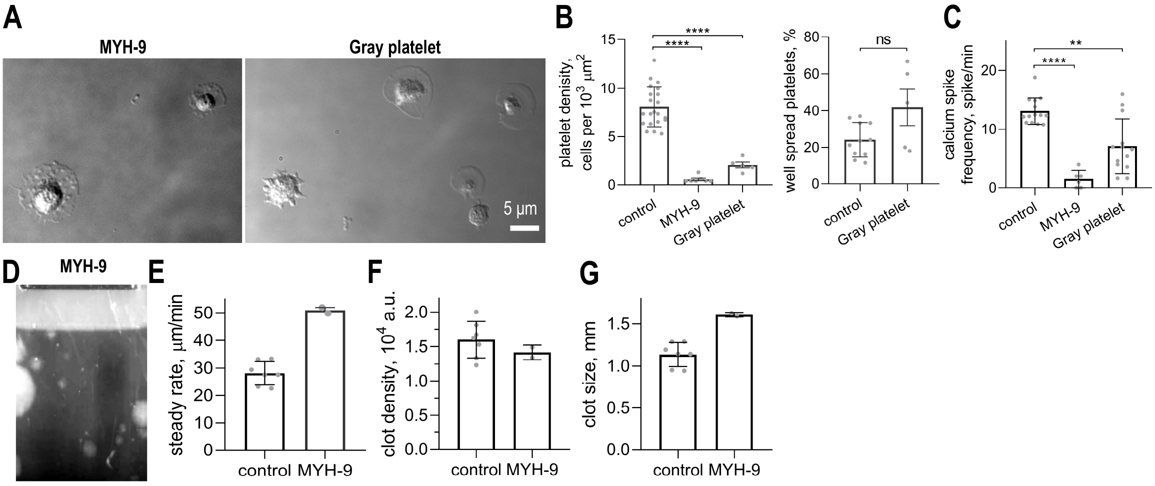


**Supplementary Figure 2. Platelet function and coagulation in MYH-9 disorder and Gray platelet syndrome patients.**

A. Representative DIC images of adherent platelets from patients with MYH-9 disorder and Gray platelet syndrome on fibrinogen-coated coverslips under flow.

B. Quantifications of platelet adhesion and spreading in MYH-9 disorder and Gray platelet syndrome samples. Left, platelet adhesion quantified as the surface density of adherent cells. Right, platelet spreading quantified as the percent of adherent platelets with visible lamellipodia. Each dot represents one field of view. Here and in panel C, bars show mean ± SD, and healthy adult controls are shown for comparison. Statistical comparisons were performed using Mann–Whitney tests; ns, not significant; ****$p < 0.0001$.

C. Calcium spike frequency in thrombin-activated platelets. Each dot represents data for one platelet. See panel B for details. Statistical comparisons were performed using Mann–Whitney tests; **$p < 0.01$; ****$p < 0.0001$.

D. Thrombodynamics image of fibrin clot in MYH-9 patient plasma at 30 min.

E. Steady-state fibrin clot growth rate in healthy adult control and MYH-9 patient plasma. Here and in panels F,G, bars show mean ± SEM; Each dot represents one independent measurement.

F. Fibrin clot density at 30 min in healthy adult control and MYH-9 patient plasma. See panel D for details.

G. Fibrin clot size at 30 min in healthy adult control and MYH-9 patient plasma. See panel D for details.

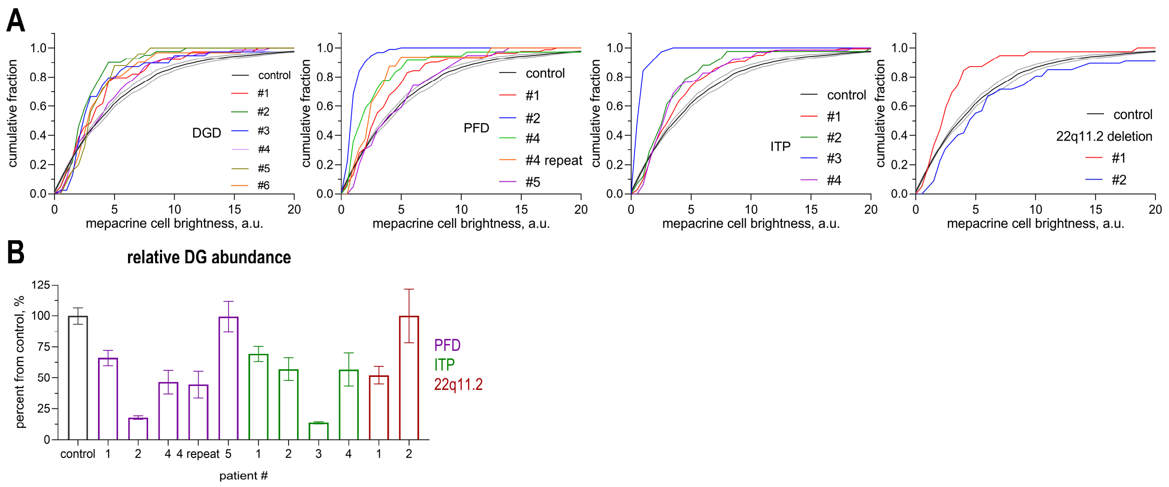


**Supplementary Figure 3. Patient-level DG content across pediatric platelet disorders.**

A. Cumulative distributions of mepacrine fluorescence in individual patients with DGD, PFD, ITP, and 22q11.2 deletion syndrome compared with healthy adult controls. Each colored curve represents one patient sample; gray curve represents averaged healthy adult controls in each graph (mean ±SEM). The repeated measurement for PFD patient #4 is shown separately. See Figure 3C for details.

B. Relative DG abundance in individual PFD, ITP, and 22q11.2 deletion syndrome patients. The first bar shows the healthy adult reference set; subsequent bars show individual patients. Values are expressed as percent of the healthy adult control value, with n = 35–140 platelets analyzed for each patient. The repeated measurement for PFD patient #4 is shown separately. Uncertainty in this estimate was calculated by bootstrap resampling of individual patient platelet fluorescence values with replacement; error bars indicate bootstrap SD.

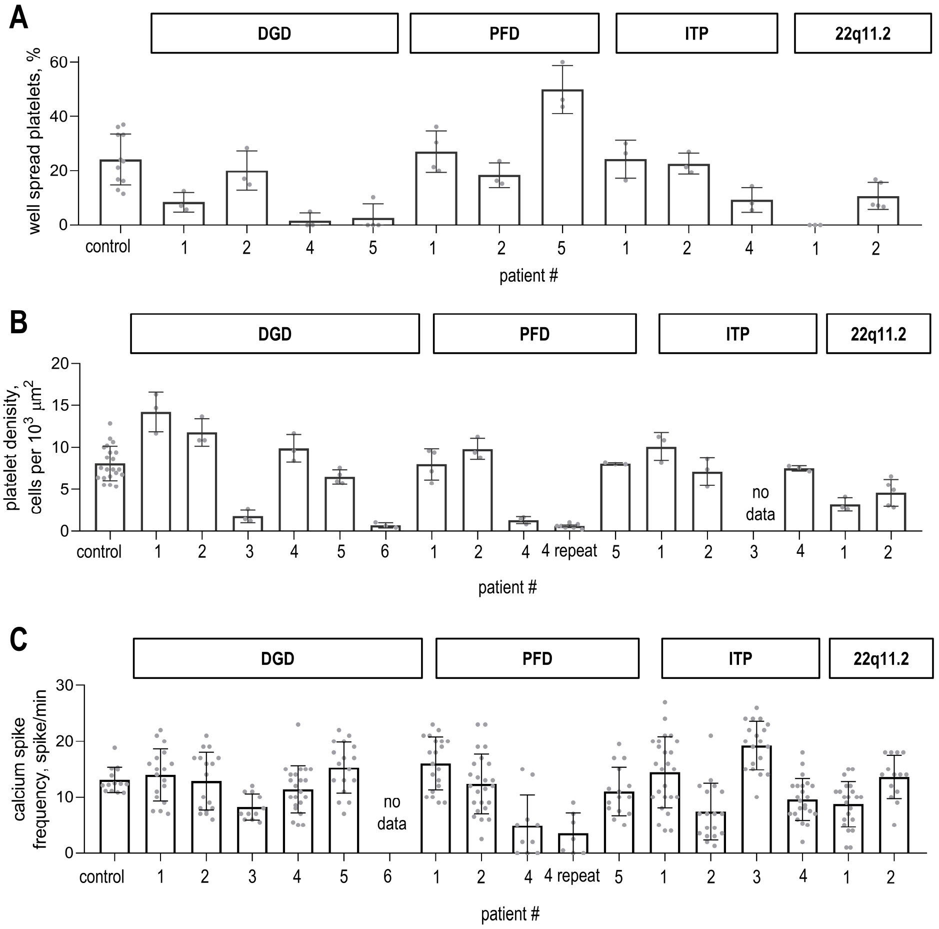


**Supplementary Figure 4. Patient-level platelet adhesion, spreading, and calcium signaling across pediatric platelet disorders.**

A. Platelet spreading in individual pediatric patients and healthy adult controls. Gray dots indicate individual fields. Here and in panels B,C, bars show mean ± SD. Patients are grouped by diagnosis, with n = 35–140 platelets analyzed for each patient.

B. Platelet adhesion in individual pediatric patients and healthy adult controls. Gray dots indicate individual fields. See panel A for details.

C. Calcium spike frequency in individual pediatric patients and healthy adult controls after thrombin stimulation. Gray dots indicate data for individual platelets. See panel A for details.

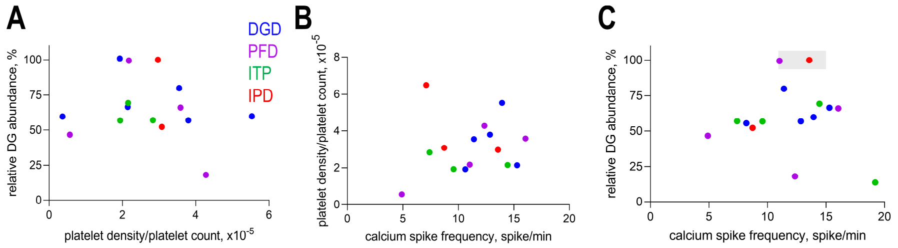


**Supplementary Figure 5. Absence of correlations between platelet functions.**

A. Relationship between relative DG abundance and platelet adhesion normalized to platelet count in blood across pediatric patient samples. Here and in panels B and C, each dot represents one patient and is color-coded by diagnostic group. Pearson correlation coefficient is -0.17.

B. Relationship between platelet adhesion normalized to platelet count in blood and calcium spike frequency across pediatric patient samples. Pearson correlation coefficient is 0.14. See panel A for details.

C. Relationship between relative DG abundance and calcium spike frequency across pediatric patient samples. Pearson correlation coefficient is 0.09. See panel A for details.

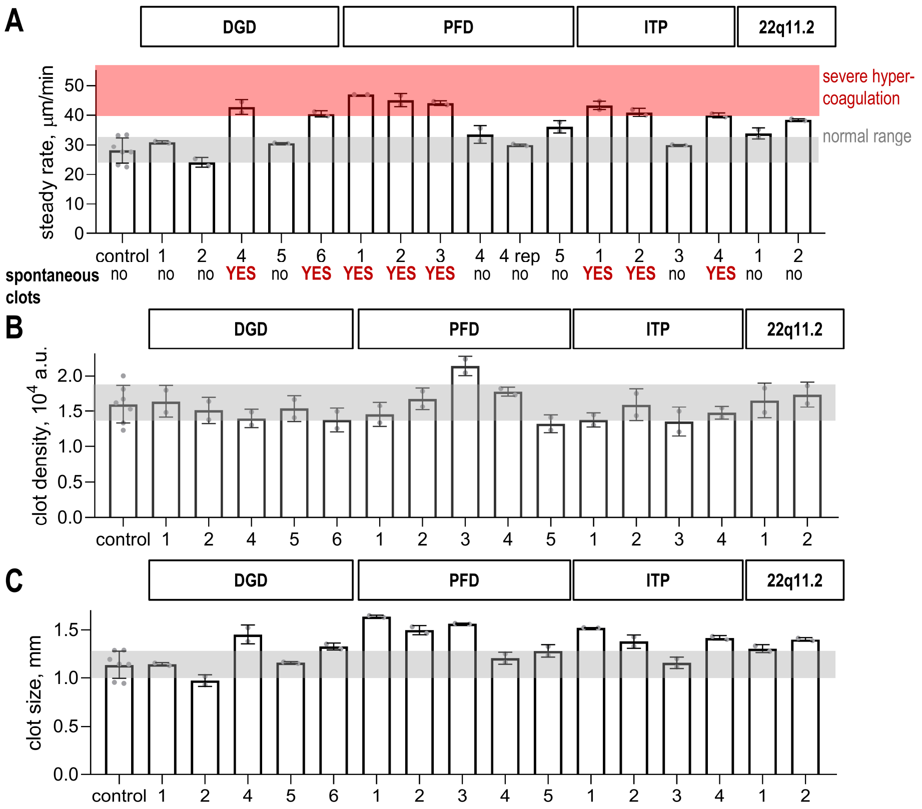


**Supplementary Figure 6. Patient-level Thrombodynamics parameters across pediatric platelet disorders.**

A. Steady-state fibrin clot growth rate in healthy adult controls and individual pediatric patient plasma samples. Here and in panels B and C, patients are grouped by diagnostic category. The gray shaded region indicates the healthy adult reference range, and the red shaded region indicates severe hypercoagulation with spontaneous clotting. Presence or absence of spontaneous clotting is indicated below each bar. Bars show mean ± SD. Gray dots represent individual measurements, with two repeats shown for each patient; for healthy adults, each dot represents the average of two repeats from one donor, N = 7.

B. Fibrin clot density at 30 min in healthy adult controls and individual pediatric patient plasma samples. See panel A for details.

C. Fibrin clot size at 30 min in healthy adult controls and individual pediatric patient plasma samples. See panel A for details.

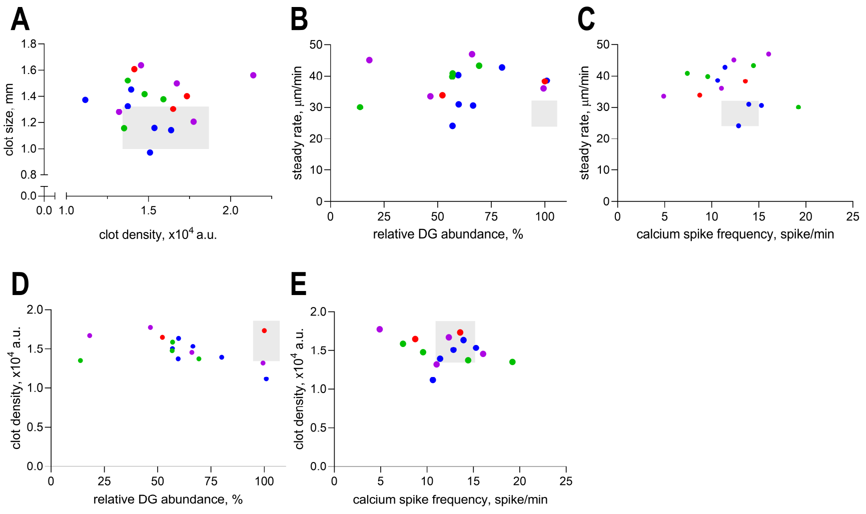


**Supplementary Figure 7. Relationships between platelet functions and plasma coagulation parameters.**

A. Relationship between clot size and clot density at 30 min across pediatric patient plasma samples. Here and in panels B-E, each dot represents one patient and is color-coded by diagnostic group. Pearson correlation coefficient is 0.08.

B. Relationship between relative DG abundance and steady-state clot growth rate across pediatric patient samples. Pearson correlation coefficient is 0.14.

C. Relationship between calcium spike frequency and steady-state clot growth rate across pediatric patient samples. Pearson correlation coefficient is -0.08.

D. Relationship between relative DG abundance and clot density across pediatric patient samples. Pearson correlation coefficient is -0.33.

E. Relationship between calcium spike frequency and clot density across pediatric patient samples. Pearson correlation coefficient is -0.31.

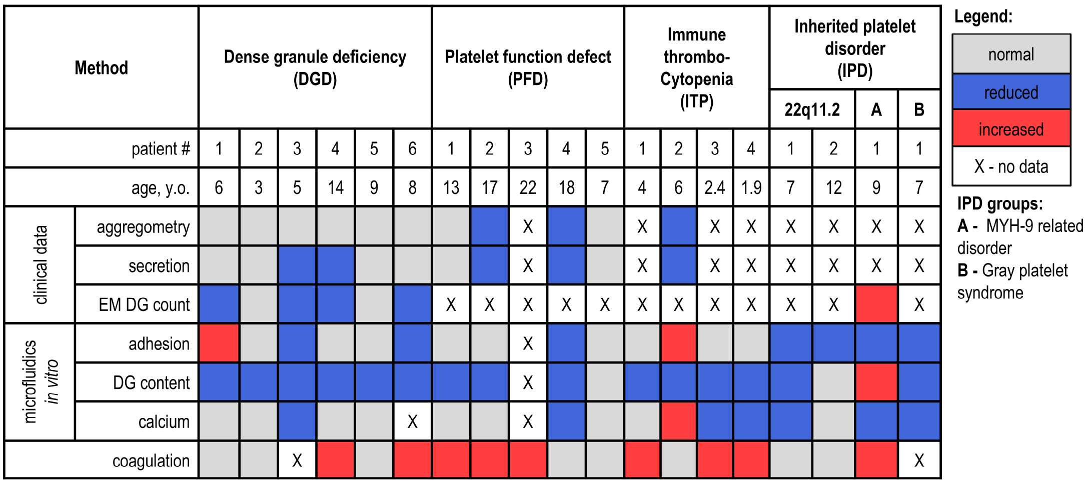


**Supplementary Figure 8. Summary table of functional phenotypes across pediatric patients.** Heatmap overview of clinical and experimental platelet parameters in patients. Rows indicate clinical testing, microfluidic in vitro assays, and plasma coagulation. Note that IPD group has subgroups.

### Supplementary Notes

#### Supplementary Note 1. Semi-automated quantification of mepacrine fluorescence intensity in individual platelets

All image analyses were performed in Fiji/ImageJ (NIH) using custom macros based on Trainable Weka Segmentation (TWS) and a pretrained classifier (“Ca classifier”). The full macro code and classifier files are provided as Supplementary Materials. DIC images and time-lapse recordings from two fluorescence channels were used for analysis: 488 nm (mepacrine), to measure fluorescence of platelet DGs, and 561 nm (Calbryte-590$^{AM}$), for platelet segmentation (Figure S1A). Frames collected before agonist addition were selected for analysis during the 8–15 min wash period (Figure 2B), after unbound mepacrine had been removed, to ensure a stable mepacrine signal.

Platelets were segmented based on the Calbryte-590$^{AM}$ channel using a custom Fiji/ImageJ macro built on the Trainable Weka Segmentation plugin[1] and a pre-trained classifier (“Ca classifier”). Segmentation steps included: 1) background subtraction using a rolling ball algorithm, 2) classifier application, 3) conversion to a binary mask, and 4) size-based selection of objects larger than 1.5 µm. Platelet ROIs were visually inspected and manually corrected when necessary (Figure S1A). A selection was considered acceptable if it was similar in size to other platelet ROIs, was not located on the image border, and did not represent overlapping cells.

The corresponding 488 nm image was background-corrected with subtraction of Gaussian blurred background, and Raw Integrated Density was measured within each platelet ROI. Raw Integrated Density values from at least three frames were averaged to obtain a single fluorescence estimate per platelet (Figure S1A).

*Validation of total cell fluorescence measurements using epifluorescence imaging.* We analyzed mepacrine fluorescence throughout z-stacks to justify using raw integrated density from a single plane as a measure of total cell fluorescence. Figures S1B and S1C show that even when a platelet appears visibly out of focus (Figure S1B), its fluorescence intensity does not change substantially (Figure S1C). This allowed us to use single-plane images from each field of view instead of z-stacks.

*Correction of total cell fluorescence.* Mepacrine is a lipophilic dye[2] and therefore can accumulate in membranes. Thus, even cells devoid of DGs can produce a fluorescence signal, which may influence Raw Integrated Density depending on cell area. To reduce this artifact, we measured the mean pixel fluorescence of empty cells, defined as platelets that had released all of their granules, as illustrated in Figure S1D. This value was averaged across 245 platelets from 17 experiments performed with blood from healthy volunteers. The resulting value was then subtracted from each pixel of the image as part of the image-processing algorithm for both patient and healthy volunteer samples, as shown in Figure S1E.
